# Impact of Thiol-Disulfide Balance on the Binding of Covid-19 Spike Protein with Angiotensin Converting Enzyme 2 Receptor

**DOI:** 10.1101/2020.05.07.083147

**Authors:** Sanchita Hati, Sudeep Bhattacharyya

## Abstract

The novel coronavirus, severe acute respiratory syndrome coronavirus 2 (SARS-CoV-2), has led to an ongoing pandemic of coronavirus disease (COVID-19), which started in 2019. This is a member of Coronaviridae family in the genus Betacoronavirus, which also includes SARS-CoV and Middle East respiratory syndrome coronavirus (MERS-CoV). The angiotensin-converting enzyme 2 (ACE2) is the functional receptor for SARS-CoV and SARS-CoV-2 to enter the host cells. In particular, the interaction of viral spike proteins with ACE2 is a critical step in the viral replication cycle. The receptor binding domain of the viral spike proteins and ACE2 have several cysteine residues. In this study, the role of thiol-disulfide balance on the interactions between SARS-CoV/CoV-2 spike proteins and ACE2 was investigated using molecular dynamic simulations. The study revealed that the binding affinity was significantly impaired when all the disulfide bonds of both ACE2 and SARS-CoV/CoV-2 spike proteins were reduced to thiol groups. The impact on the binding affinity was less severe when the disulfide bridges of only one of the binding partners were reduced to thiols. This computational finding provides a molecular basis for the severity of COVID-19 infection due to the oxidative stress.

## 1. INTRODUCTION

The novel coronavirus known as severe acute respiratory syndrome coronavirus 2 (SARS-CoV-2) or simply COVID-19 is the seventh member of the coronavirus family.^1^ The other two viruses in this family that infect humans are severe acute respiratory syndrome coronavirus (SARS-CoV) and Middle East respiratory syndrome coronavirus (MERS-CoV). These are positive-sense, single-strand enveloped RNA viruses. The coronavirus particles contain four main structural proteins: the spike, membrane, envelope, and nucleocapsid.^1-2^ The spike protein protrudes from the envelope of the virion and consists of two subunits; a receptor-binding domain (RBD) that interacts with the receptor proteins of host cells and a second subunit that facilitates fusion of the viral membrane into the host cell membrane. Recent studies showed that RBD of spike proteins of SARS-CoV and SARS-CoV-2 interact with angiotensin-converting enzyme 2 (ACE2). ACE2 belongs to the membrane-bound carboxydipeptidase family. It is attached to the outer surfaces of cells and is widely distributed in the human body. In particular, higher expression of ACE2 is observed in organs such as small intestine, colon, kidney, and heart, while ACE2 expression is comparatively lower in liver and lungs.^3-4^

The role of oxidative stress on the binding of viral proteins on the host cell surface receptors is a relatively underexplored area of biomedical research.^5-10^ Previous studies have indicated that the entry of viral glycoprotein is impacted by thiol-disulfide balance on the cell surface.^5, 7-9, 11-12^ Any perturbations in the thiol-disulfide equilibrium has also been found to deter the entry of viruses into their target cells.^5^ The first step of the viral entry involves binding of the viral envelop protein onto a cellular receptor. This is followed by endocytosis, after which conformational changes of the viral protein helps the induction of the viral protein into the endosomal membrane, finally releasing the viral content into the cell. These conformational changes are mediated by pH changes as well as the conversion of disulfide to thiol group of the viral spike protein.^7^ Several cell surface oxidoreductases^9^ regulate the thiol-disulfide exchange, responsible for conformational changes of viral proteins needed for virus entry into host cells.

In the backdrop of significant mortality rate for SARS-CoV-2 (hereinafter referred to as CoV-2) infection, it is important to know if the thiol-disulfide balance plays any role on the binding of the spike glycoprotein on to the host cell receptor protein ACE2. A recent study with the spike glycoprotein of SARS-CoV (hereinafter referred to as CoV) has exhibited a complete redox insensitivity;^7^ despite the reduction of all disulfide bridges of CoV to thiols, its binding to ACE2 remained unchanged.^7^ However, this study did not probe the redox sensitivity of ACE2 receptor. Thus, in the present study, we computationally investigated the redox state of both partners (ACE2 and CoV/CoV-2) on their binding affinities. The structure of CoV^13^ and CoV-2^14-15^ complexed with ACE2 are known and the noncovalent interactions at the protein-protein interface^16^ have been reported recently. Using these reported structures, molecular dynamics simulations and electrostatic field calculations were performed to explore the impact of thiol-disulfide balance on CoV/CoV-2 and ACE2 binding affinities. The structural and dynamical changes due to the change in the redox states of cysteines in the interacting proteins were analyzed and their effects on binding free energies were studied.

## 2. RESULTS AND DISCUSSION

The molecular basis of the binding of spike proteins to ACE2 is known from X-ray crystallographic (SARS-CoV)^13^ and cryo-electron microscopic (SARS-CoV-2)^16^ studies. The sequence alignment of CoV and CoV-2 spike proteins showed high sequence identity (>75 %) indicating that their binding to ACE2 receptors will be similar (Fig. 1). In both bound structures, the RBD of CoV and CoV-2 is found to be complexed with ACE2 (Fig. 2). Both ACE2 and CoV-2 possess four disulfide bridges, whereas CoV subunit has only two disulfide linkages (Table 1 and Fig. 2a and 2b). Two large helices of ACE2 form a curved surface (Fig. 2, illustrated by the dashed curved line) that interacts with the concave region of CoV or CoV-2 subunit.

**Table 1.**
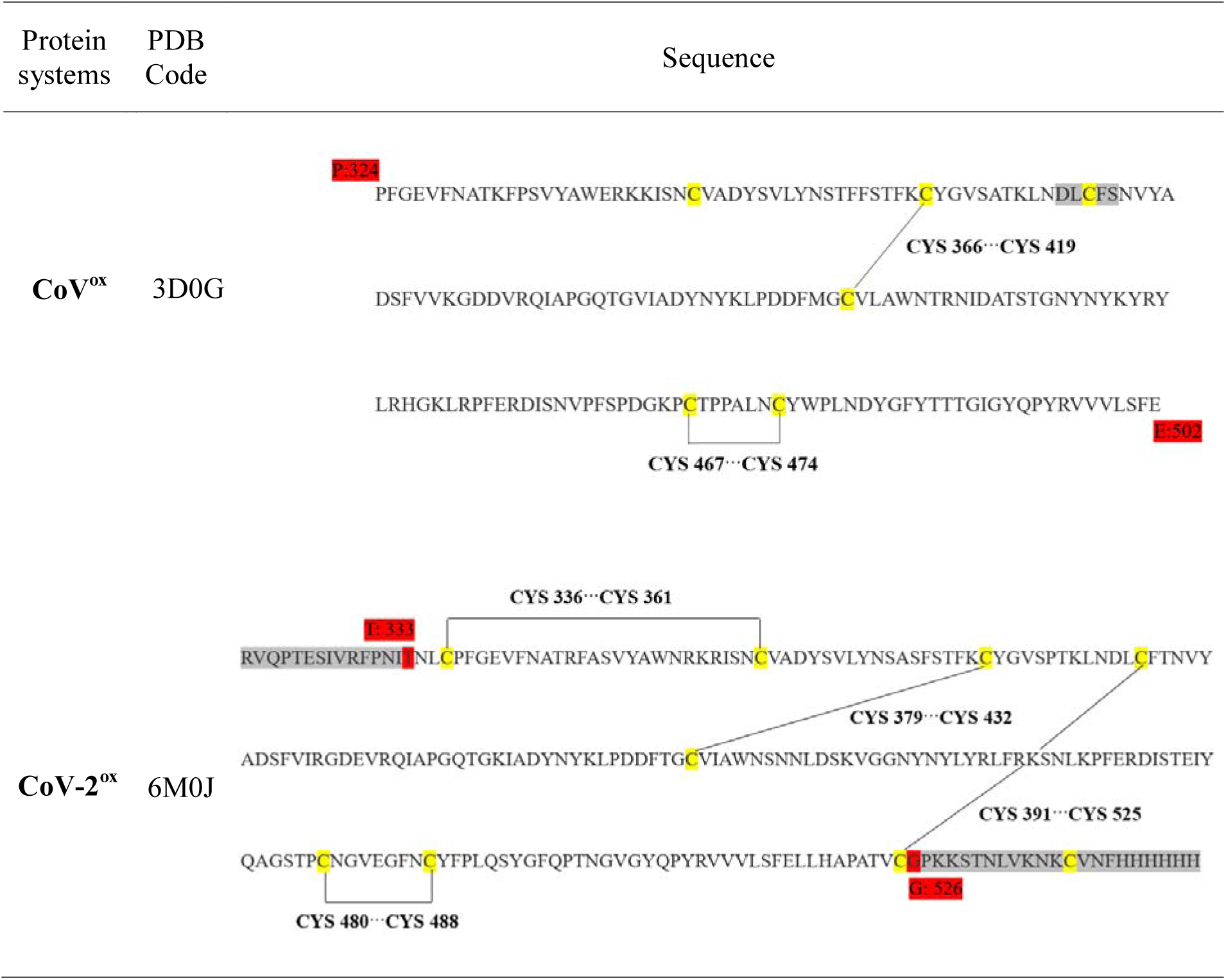
Sequences of the receptor-binding domain of SARS-CoV and SARS-CoV2 proteins. The cysteine residues re highlighted in yellow and disulfide bonds are shown using black solid lines. The start and end residues are numbered and highlighted in red. The gray highlighted residues are missing in the crystal structure.

**Figure 1.**
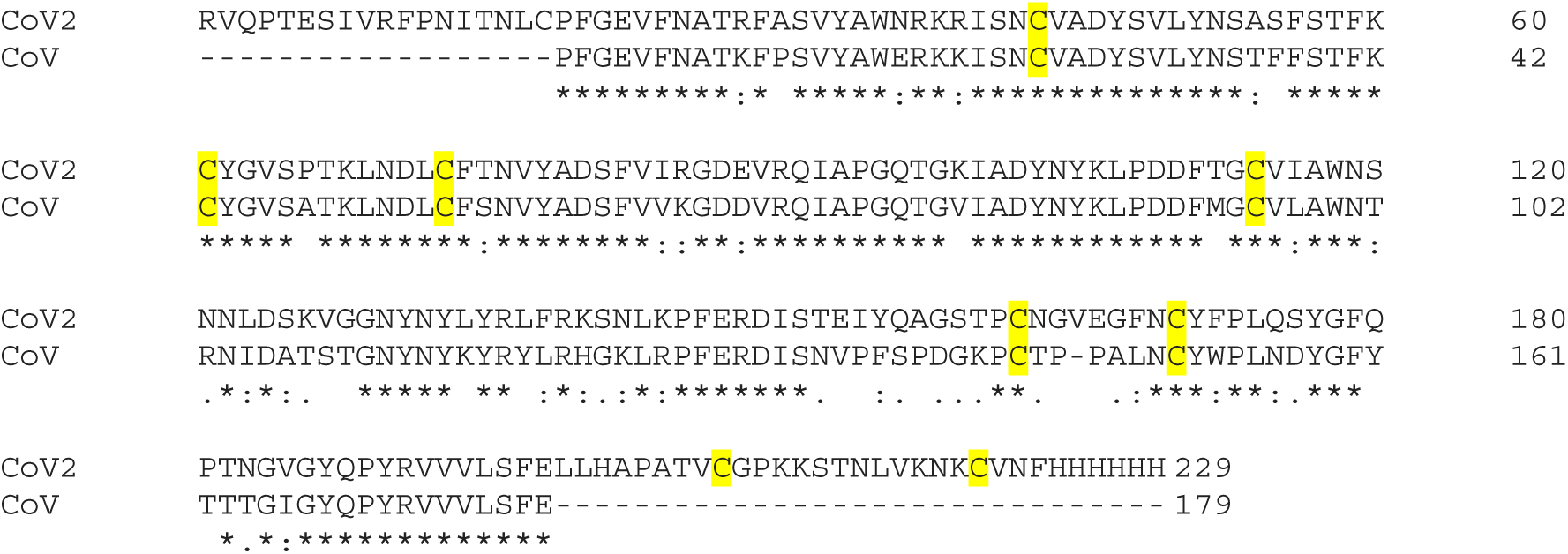
Sequence alignment (generated by Clustal Omega^35^) between the receptor-binding domain of SARS-CoV and SARS-CoV2 proteins. The “*” represents the identical residues, “:” represents similar residues, and gap represents dissimilar residues. The cysteine residues are highlighted in yellow.

**Figure. 2.**
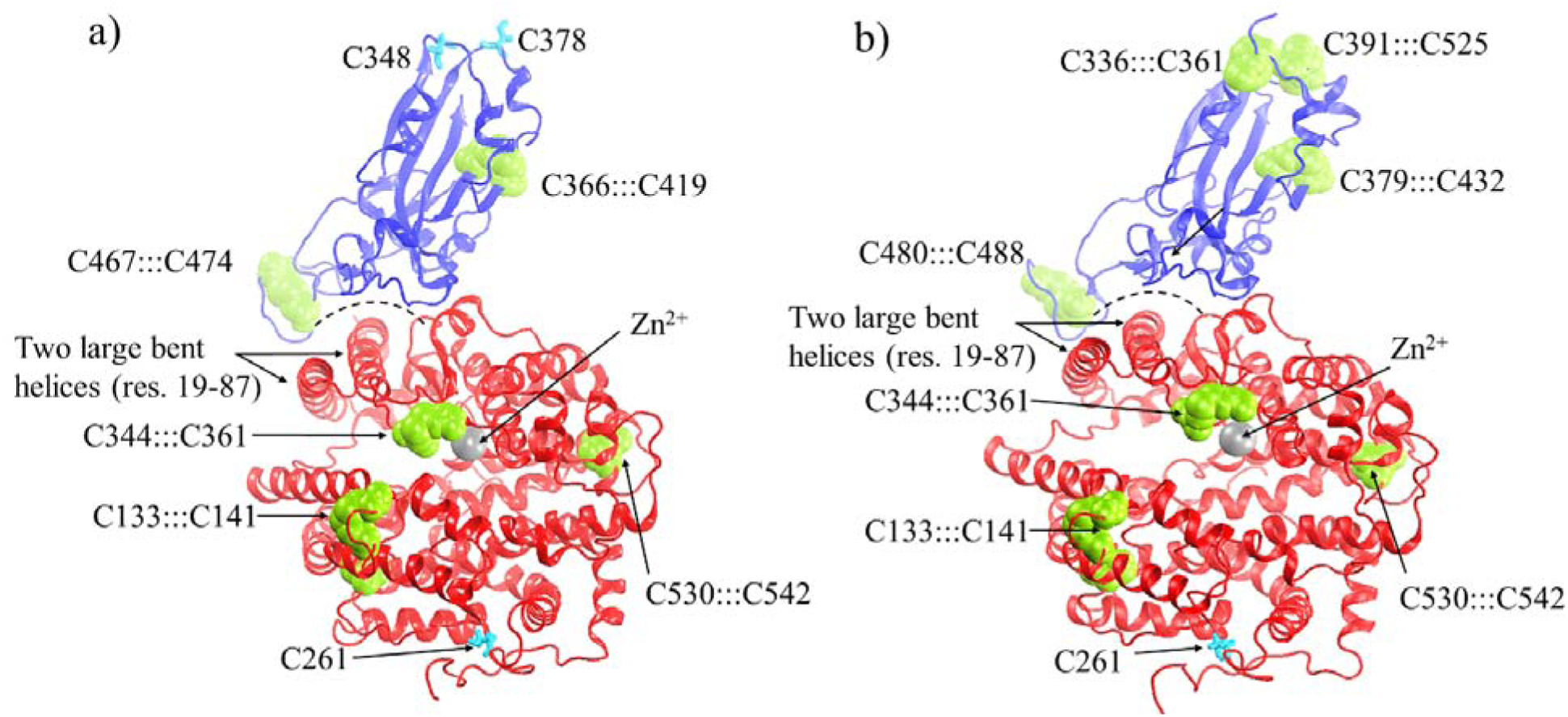
Structures of protein complexes of a) SARS-CoV…ACE2 and b) SARS-CoV-2…ACE2. All the disulfide bridges between cysteine residues are shown in green vdW spheres and thiol groups in cyan licorice.

### Structural change along the trajectory

In all cases, the simulation started with an equilibrated structure, which was obtained after minimizing the neutralized solvated protein complex built from the experimentally determined structures. The evolution of the protein structure along the MD trajectory was monitored by calculating the root-mean-square deviation (RMSD) of each structure from the starting structure as a frame of reference following standard procedure.^17^ Briefly, during the MD simulation, the protein coordinates were recorded for every 10 ps interval and a root-mean-square-deviation (RMSD) of each frame was calculated from the average root-mean-square displacement of backbone Cα atoms with respect to the initial structure. Then, the RMSD values, averaged over conformations stored during 1 ns time, were plotted against the simulated time (Fig. 3). Compared to the starting structure, only a moderate backbone fluctuation was noted in all protein complexes during 20 ns simulations and the maximum of RMSD was in the range of 2.0-3.8 Å (Table 2). The evolution was smooth and its stability was demonstrated by the standard deviation of the computed RMSDs, which was less than 0.3 Å. Taken together, results of the simulations showed no unexpected structural deformation of the SAR-CoV/CoV-2…ACE2 complex in both, reduced (thiol groups) and oxidized (disulfide bonds) states (Table 2).

**Table 2.**
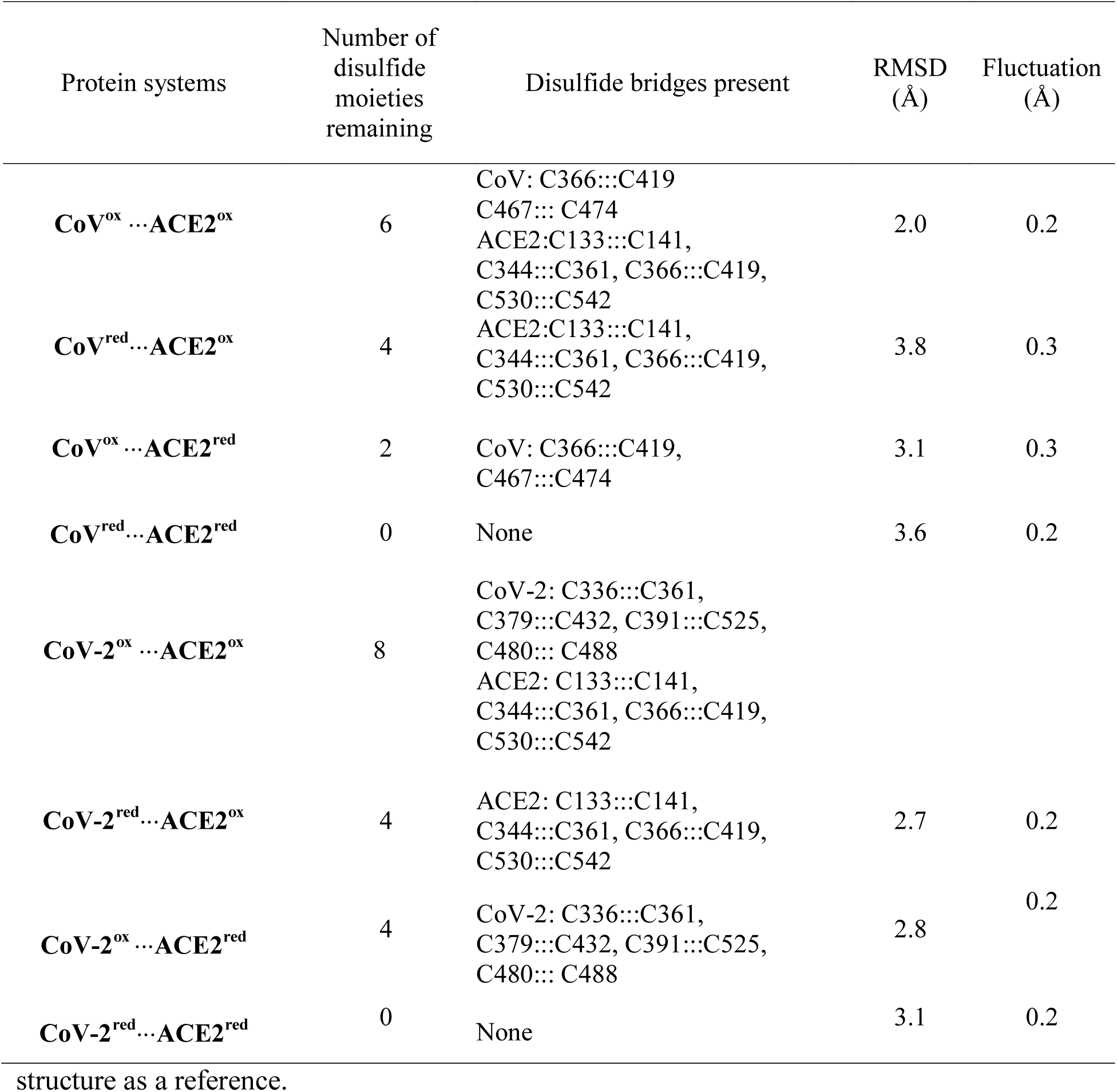
Evolution of the structure of the protein complexes for the oxidized forms and other variants observed during 20 ns molecular dynamics simulations. For each protein system, the ‘RMSD’ column contains the average of the last 5 ns RMSD, and the ‘Fluctuation’ column represents the standard deviation during the same duration computed based on the starting

| Protein systems | Number of disulfide moieties remaining | Disulfide bridges present | RMSD (Å) | Fluctuation (Å) |
| --- | --- | --- | --- | --- |
| **CoV<sup>ox</sup> ...ACE2<sup>ox</sup>** | 6 | CoV: C366:::C419<br>C467::: C474<br>ACE2:C133:::C141,<br>C344:::C361, C366:::C419,<br>C530:::C542 | 2.0 | 0.2 |
| **CoV<sup>red</sup> ...ACE2<sup>ox</sup>** | 4 | ACE2:C133:::C141,<br>C344:::C361, C366:::C419,<br>C530:::C542 | 3.8 | 0.3 |
| **CoV<sup>ox</sup> ...ACE2<sup>red</sup>** | 2 | CoV: C366:::C419,<br>C467:::C474 | 3.1 | 0.3 |
| **CoV<sup>red</sup> ...ACE2<sup>red</sup>** | 0 | None | 3.6 | 0.2 |
| **CoV-2<sup>ox</sup> ...ACE2<sup>ox</sup>** | 8 | CoV-2: C336:::C361,<br>C379:::C432, C391:::C525,<br>C480::: C488<br>ACE2: C133:::C141,<br>C344:::C361, C366:::C419,<br>C530:::C542 |  |  |
| **CoV-2<sup>red</sup> ...ACE2<sup>ox</sup>** | 4 | ACE2: C133:::C141,<br>C344:::C361, C366:::C419,<br>C530:::C542 | 2.7 | 0.2 |
| **CoV-2<sup>ox</sup> ...ACE2<sup>red</sup>** | 4 | CoV-2: C336:::C361,<br>C379:::C432, C391:::C525,<br>C480::: C488 | 2.8 | 0.2 |
| **CoV-2<sup>red</sup> ...ACE2<sup>red</sup>** | 0 | None | 3.1 | 0.2 |

structure as a reference.

**Figure 3.**
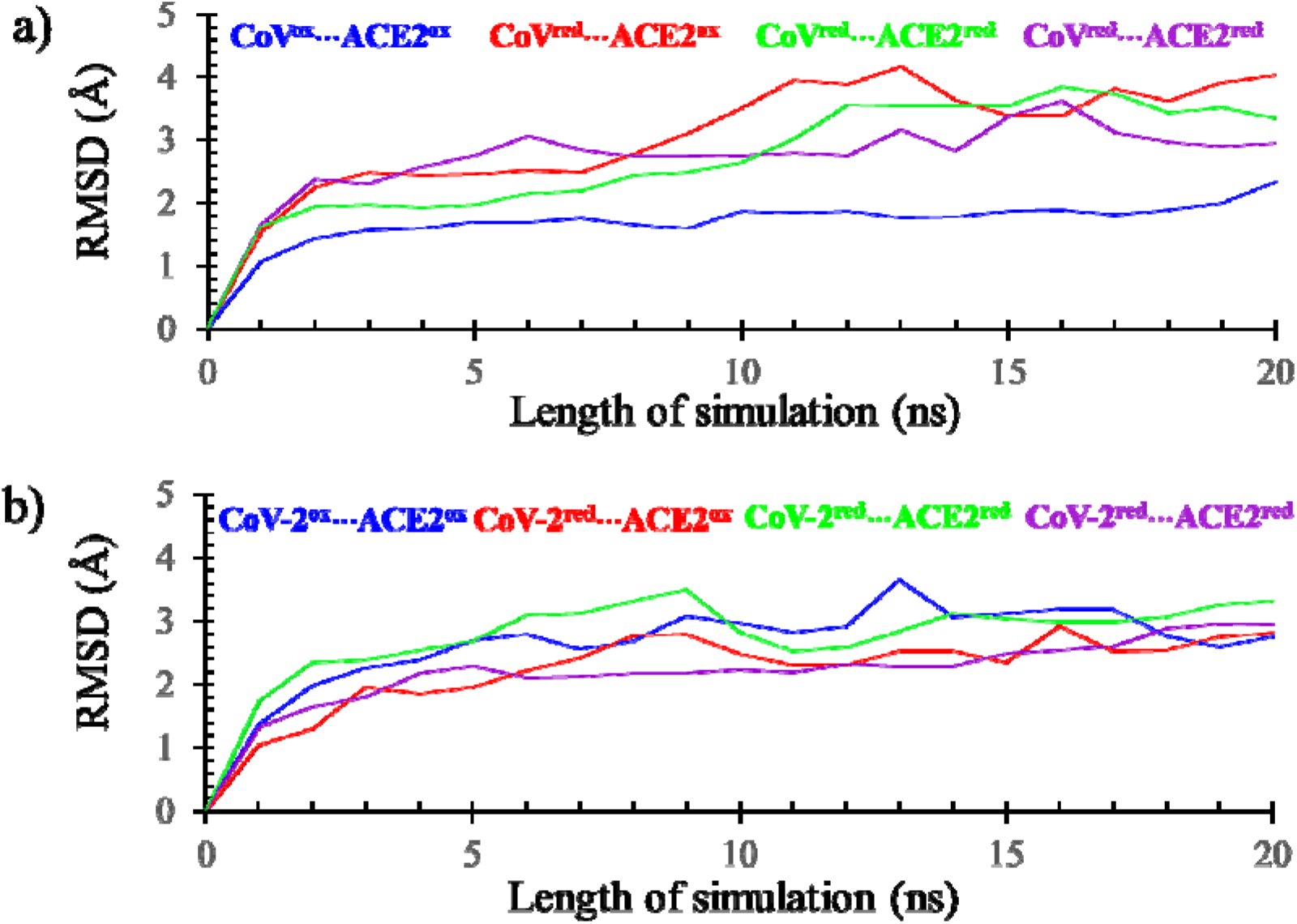
RMSDs, averaged over 1 ns MD simulation, of the protein complexes of ACE2 and SARS-CoV or SARS-CoV-2. Disulfide-containing proteins are referred as oxidized with a shorthand notation of ‘ox’, while thiol variants are denoted with ‘red’ notation.

### Thermal fluctuation due to cleavage of disulfide bridges

The MD simulation results indicated that the flexibility of different structural elements of the interacting proteins altered upon the cleavage of disulfide bridges (oxidized state) producing sulfydryl (thiol) groups (reduced state). The impact of the reduction of the disulfide bridges was probed by calculating per-residue RMSD of each complex from the trajectories of MD simulations. Changes in the backbone flexibility due to complete reduction of all disulfides to thiols of both interacting proteins are shown in Figs. 4a and 4b, where the backbone is color-coded to indicate the magnitude of thermal fluctuations. Larger and smaller backbone fluctuations are shown in red and blue, respectively, whereas the medium fluctuation regions are shown in green. The backbone fluctuation analysis demonstrated that the cleavage of disulfide bridges indeed affected the flexibility of the local regions as they became more dynamic (Fig. 4b). A significant alteration in backbone flexibility was noticed at the interface of CoV and ACE2 subunits when both proteins are in the fully reduced states i.e. all disulfides are changed to sulfydryl groups. As it is evident from the Fig. 4b, the helix (residues 52-64) of ACE2 became very mobile, due to the rupture of the nearby C344:::C361 disulfide. The second region belongs to the flexible loop consisting residues 463-474 of CoV subunit (residues 478-489 of CoV-2 subunit), which interacts with the N-terminal segment of the ACE2. The cleavage of the disulfides C467:::C474 in CoV and C480:::C488 in CoV-2 resulted in higher thermal fluctuation.

**Figure 4.**
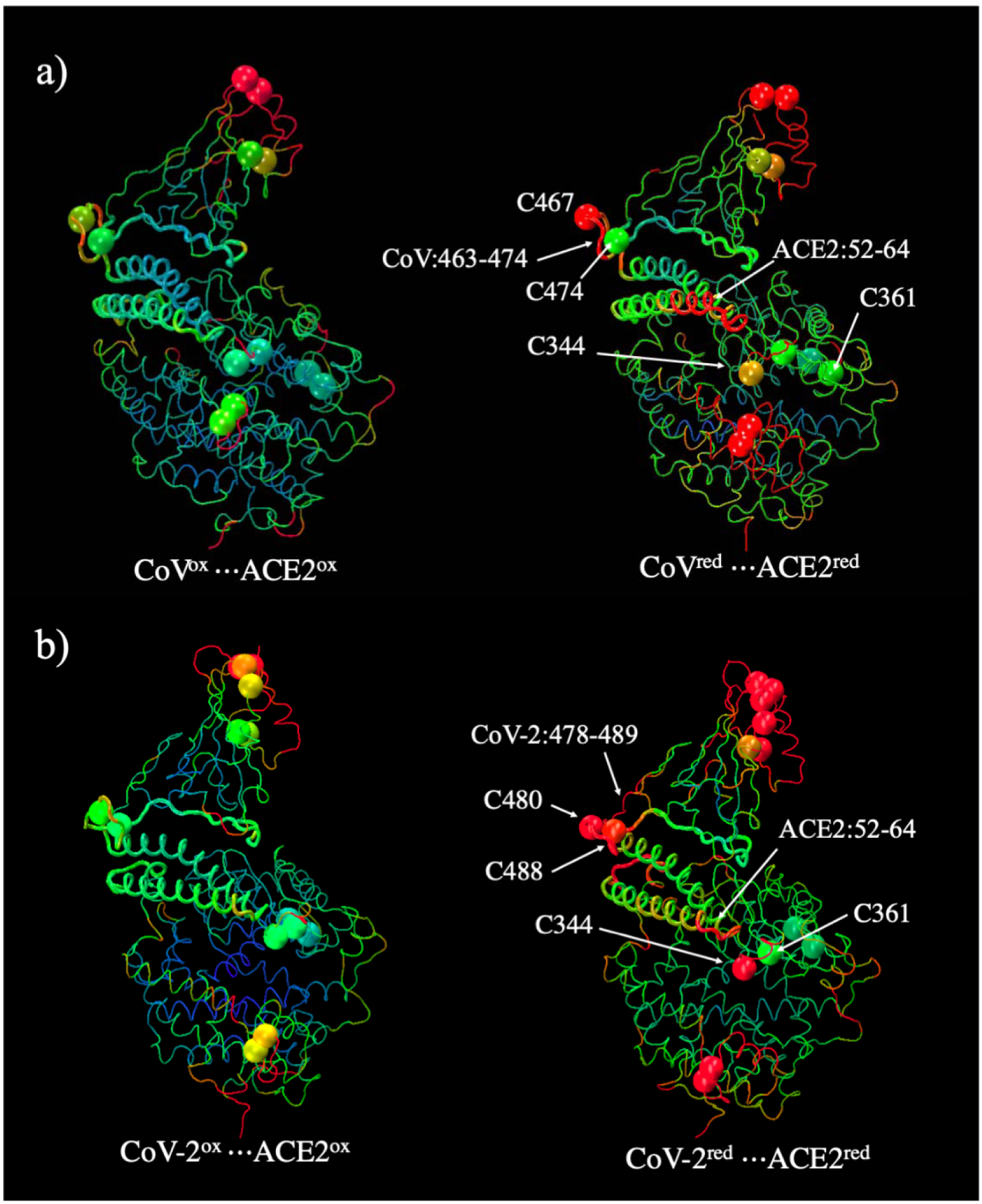
Change in backbone fluctuations in the oxidized and reduced protein complexes: a) SARS-CoV…ACE2 and b) SARS-CoV-2…ACE2. The cysteine residues, for each protein systems (labeled in Fig. 2) are highlighted as vdW spheres. The backbone as well as the cysteine residues are color coded; large fluctuations are shown in red and small fluctuations are in blue. The medium-scale fluctuations are shown in green.

### Binding study

The computed Gibbs binding free energy (Δ_bind_*G*^o^) of protein complexes (Table 3) demonstrated that the binding of CoV or CoV-2 with ACE2 occurs because the attractive electrostatic interactions between the individual subunits prevail over the desolvation due to the complexation. The complex formation results in desolvation from proteins’ surfaces at the protein-protein interface, thereby producing a positive ΔΔ_solv_*G*_corr_ for all protein systems (Table 3). For formation of a stable protein-protein complex, this energy is needed to be counterbalanced by the negative electrostatic interactions, Δ _Coul_*G*, between the interacting proteins.

**Table 3.**
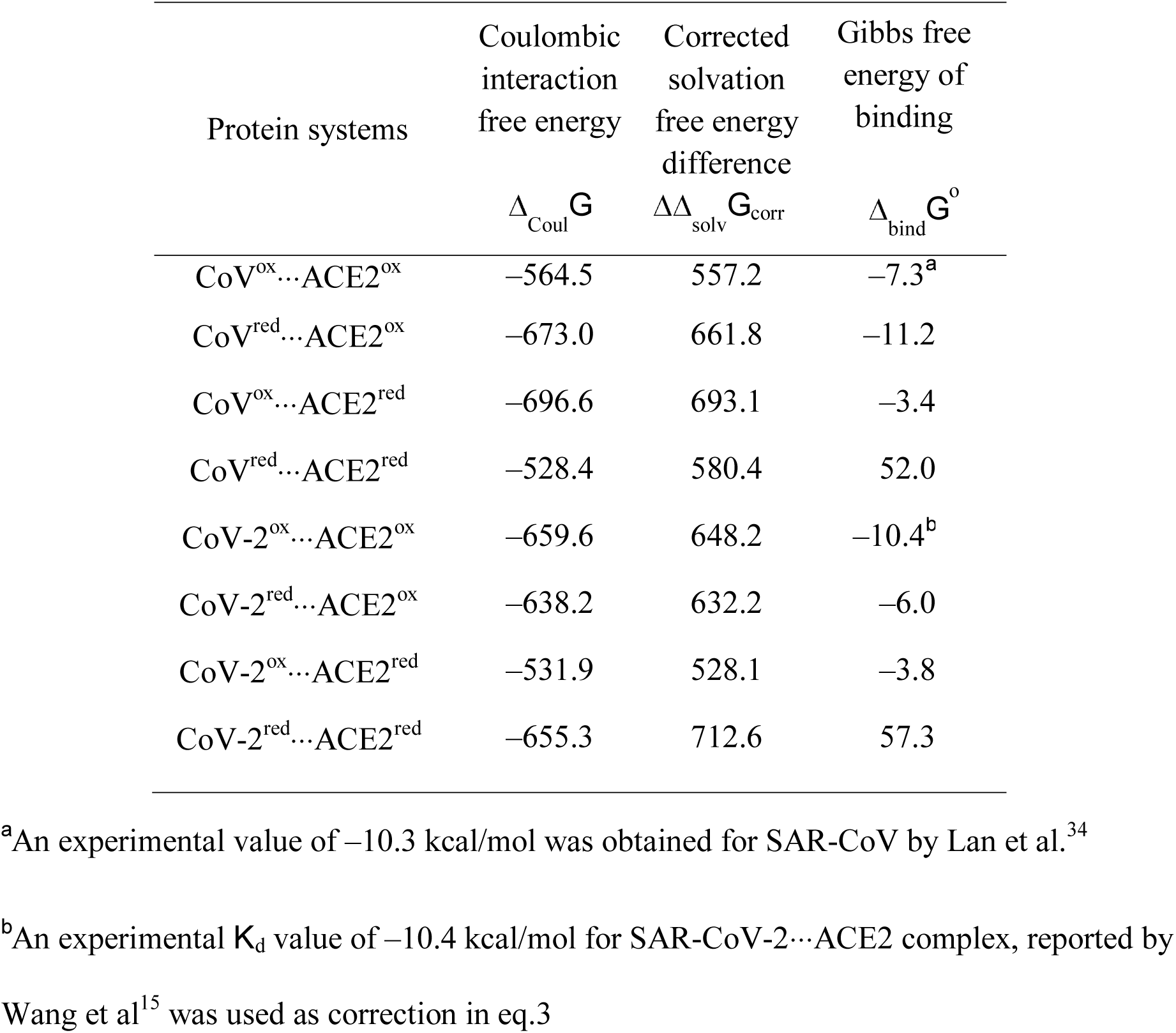
Various components of the Gibbs free energy of binding calculated by implicit solvation and APBS. All energies are expressed in kcal/mol. An estimated uncertainty of 1-3 kcal/mol was determined for the computed Gibbs free energy.

| Protein systems | Coulombic<br>interaction<br>free energy<br>$\Delta_{\text{Coul}} G$ | Corrected<br>solvation<br>free energy<br>difference<br>$\Delta\Delta_{\text{solv}} G_{\text{corr}}$ | Gibbs free<br>energy of<br>binding<br>$\Delta_{\text{bind}} G^o$ |
| --- | --- | --- | --- |
| CoV<sup>ox</sup>...ACE2<sup>ox</sup> | -564.5 | 557.2 | -7.3<sup>a</sup> |
| CoV<sup>red</sup>...ACE2<sup>ox</sup> | -673.0 | 661.8 | -11.2 |
| CoV<sup>ox</sup>...ACE2<sup>red</sup> | -696.6 | 693.1 | -3.4 |
| CoV<sup>red</sup>...ACE2<sup>red</sup> | -528.4 | 580.4 | 52.0 |
| CoV-2<sup>ox</sup>...ACE2<sup>ox</sup> | -659.6 | 648.2 | -10.4<sup>b</sup> |
| CoV-2<sup>red</sup>...ACE2<sup>ox</sup> | -638.2 | 632.2 | -6.0 |
| CoV-2<sup>ox</sup>...ACE2<sup>red</sup> | -531.9 | 528.1 | -3.8 |
| CoV-2<sup>red</sup>...ACE2<sup>red</sup> | -655.3 | 712.6 | 57.3 |

<sup>a</sup>An experimental value of -10.3 kcal/mol was obtained for SAR-CoV by Lan et al.<sup>34</sup>
<sup>b</sup>An experimental $K_d$ value of -10.4 kcal/mol for SAR-CoV-2...ACE2 complex, reported by Wang et al<sup>15</sup> was used as correction in eq.3

When the disulfides of the RBD of CoV and CoV-2 were reduced to thiols, the impact of reduction on the binding of these two proteins with ACE2 was different. The computed Δ_bind_*G*^o^ for CoV^ox^ …ACE2^ox^ and CoV^red^ …ACE2^ox^ are −7.3 kcal/mol and −11.2 kcal/mol, respectively (Table 3) indicating a slightly tighter binding of the reduced CoV RBD to ACE2. The experimentally known *K*_d_ value for CoV^ox^…ACE2^ox^ system is 325 nM,^14^ which corresponds to a Δ_bind_*G*^o^ value of −8.9 kcal/mol. Therefore, the computationally determined Δ_bind_*G*^o^ for CoV^ox^ …ACE2^ox^ binding is comparable to the experimental value. Although, there is no experimentally determined *K*_d_ available for CoV^red^ …ACE2^ox^ system, the binding assay with CoV^red^ RBD found to have no redox sensitivity for its binding to ACE2.^7^ The small favorability of CoV^red^…ACE2^ox^ binding, as observed from the computation, indicates that the reduction of CoV did not have much impact on its binding affinity for ACE2 and could explain the experimentally observed redox insensitivity.s^7^ In the case of CoV-2, its binding with ACE2 became less favorable when all the disulfides of RBD were reduced to thiols. The computed Gibbs binding free energies was increased by ∼ 4.5 kcal/mol for the reduced CoV-2; Δ_bind_*G*^o^ of CoV-2^ox^…ACE2^ox^ and CoV-2^red^…ACE2^ox^ are −10.4 kcal/mol and −6.0 kcal/mol, respectively.

In contrast, the reduction of disulfides of ACE2 impaired the binding significantly for both CoV and CoV-2 proteins. The Gibbs binding free energies of viral spike proteins CoV and CoV-2 with ACE2^red^, namely, CoV^ox^…ACE2^red^ and CoV-2^ox^…ACE2^red^ are –3.4 kcal/mol and –3.8 kcal/mol, respectively. These Δ_bind_*G*^o^ values are ∼ 4 - 6 kcal/mol more positive than what were observed for the oxidized from of ACE2 (i.e. CoV^ox^…ACE2^ox^ and CoV-2^ox^…ACE2^ox^ in Table 3). However, when all disulfides in CoV/CoV-2 as well as ACE2 were reduced to thiols, the binding became thermodynamically unfavorable. In both cases, the binding free energies have positive values; Δ_bind_*G*^o^ of CoV^red^…ACE2^red^ and CoV-2^red^…ACE2^red^ are 52.0 kcal/mol and 57.3 kcal/mol, respectively. These results indicate that the binding will be severely impacted, when the disulfides of both interacting proteins are converted to thiols. This finding is potentially significant as it indicates that the cleavage of disulfide bridges in ACE2 has significant destabilizing effect on the spike protein binding.

### Molecular basis of the impaired binding

The conjoined two helices of ACE2 (residues 19-53 and residues 56-94) form a complementary shape (Fig. 5) that fits into the concave-shaped loop-β-sheet-loop motif (residues 460-490 of CoV and residues 475-505 of CoV-2) of RBD of viral spike proteins. The loss of disulfide bridges in ACE2 made a strong impact on the junction of the two conjoined helices (Fig. 5, CoV^red^…ACE2^red^). This region is close to the disulfide bridge C344:::C361 and its cleavage has significant impact on the shape complementarity at the protein-protein interface. Similarly, the loss of C467:::C474 disulfide in CoV (C480:::C488 in CoV-2) resulted in a significant conformational change in the loop region, which was displaced away from the protein-protein interface by ∼6 Å indicating a reduced binding interactions as confirmed by the binding free energy calculations. This conformational change as well as the alteration in the backbone flexibility (Fig. 4) must have resulted in impaired binding of CoV/CoV-2 with ACE2.

**Figure 5.**
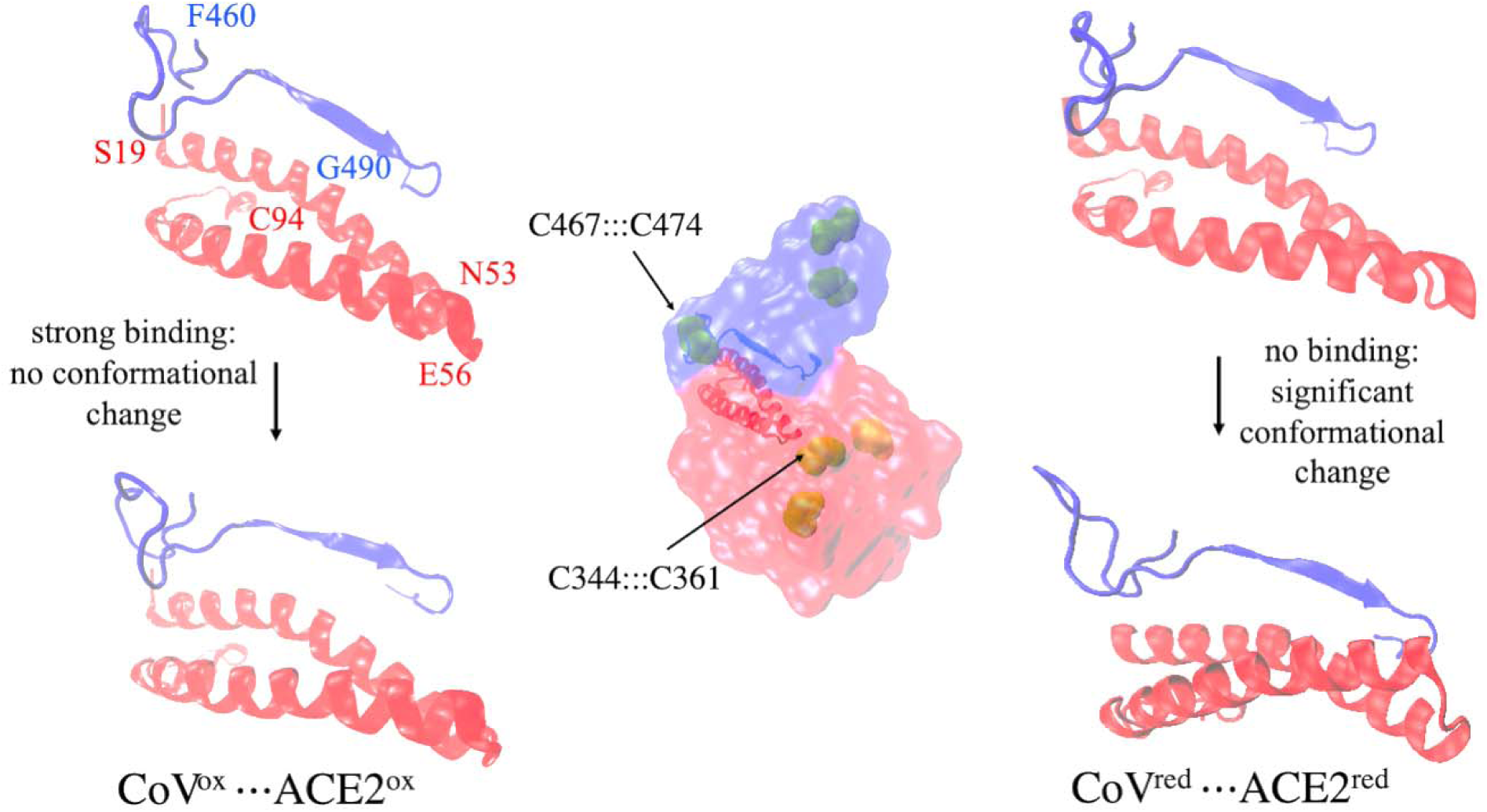
A comparison of the conformational change at the protein-protein interface in CoV^ox^ …ACE2^ox^ (left) and CoV^red^ …ACE2^red^ (right). CoV and ACE2 subunits in the complex are shown in blue and red colors, respectively, at the center. The structural motif containing the two helices of ACE2 and a beta sheet of the CoV (or CoV-2) was monitored before (top) and after (bottom) 20 ns MD simulation. The left and right panels show the difference in conformational changes in the oxidized form (CoV^ox^ …ACE2^ox^) and reduced form (CoV^red^ …ACE2^red^), respectively.

## 3. CONCLUSIONS

The study found that the reduction of all disulfides into sulfydryl groups completely impairs the binding of SARS-CoV/CoV-2 spike protein to ACE2. This is evident from the positive Gibbs energy of binding (Δ_bind_*G*^o^) obtained for both CoV^ox^…ACE2^ox^ and CoV-2^ox^…ACE2^ox^ complexes. When the disulfides of only ACE2 were reduced to sulfydryl groups, the binding becomes weaker, as the Δ_bind_*G*^o^ becomes less negative by 4 - 6 kcal/mol. On the other hand, reduction of disulfides of the RBD of CoV-2 has comparatively less effect on Δ_bind_*G*^o^ and CoV does not impact the binding with ACE2. This finding is consistent with the observed redox insensitivity of the binding between CoV and ACE2.^7^

The redox environment of cell surface receptors is regulated by the thiol-disulfide equilibrium in the extracellular region.^12, 18^ This is maintained by glutathione transporters,^19^ a number of oxidoreductases^12^ including protein disulfide isomerase,^8^ and several redox switches.^12^ Under oxidative stress, the extracellular environment becomes oxidation-prone resulting more disulfide formation on protein surfaces.^12^ Therefore, under severe oxidative stress, the cell surface receptor ACE2 and RBD of the intruding viral spike protein are likely to be present in its oxidized form having predominantly disulfide linkages. This computational study shows that under oxidative stress, the lack of reducing environment would result in significantly favorable binding of the viral protein on the cell surface ACE2. In terms of energetics, this computational study demonstrates that the oxidized form of proteins with disulfide bridges would cause a 50 kcal/mol of decrease in Gibbs binding free energy. Furthermore, ACE2, which the viral spike proteins latch on to, is known to be a key player in the remedial of oxidative stress.^20^ Binding of the viral protein will prevent the key catalytic function of ACE2 of converting angiotensin 2 (a strong activator of oxidative stress) to angiotensin 1-7 thereby creating a vicious circle of enhanced viral attack. In summary, the present study demonstrates that the absence of or reduced oxidative stress would have a significant beneficial effect during early stage of viral infection by preventing viral protein binding on the host cells.

## 4. METHODOLOGY

### Computational Setup

Setting up of protein systems and all structural manipulations were carried out using Visual Molecular Dynamics (VMD).^21^ Disulfide groups were modified to thiols during setting up of structures using standard VMD scripts. Molecular optimization and dynamics (MD) simulations were carried out using Nanoscale Molecular Dynamics (NAMD) package using CHARMM36 force field.^22-26^ During MD simulations, electrostatic energy calculations were carried out using particle mesh Ewald method.^27^ Backbone root-mean-square-deviation (RMSD) calculations were performed using VMD. Protein-protein interactions were studied using Adaptive-Basis Poisson Boltzmann Solver (APBS).^28^ Electrostatic field calculations were performed using PDB2PQR program suit.^29^

### Molecular Dynamics Simulations

All simulations were performed using the structure of ACE2 bound SARS-CoV (PDB entry: 3D0G)^13^ and SCAR-CoV-2 (PDB entry: 6M0J)^15^. In all simulations, setup of protein complexes systems was carried out following protocols used previous studies from this lab.^17, 30^ Briefly, hydrogens were added using the HBUILD module of CHARMM. Ionic amino acid residues were maintained in a protonation state corresponding to pH 7. The protonation state of histidine residues was determined by computing the p*K*a using Propka option of PDB2PQR. The protein structures were explicitly solvated with water of TIP3P model and neutralized by adding with 24 sodium atoms. The prepared solvated protein complexes were of dimensions 92 Å × 132 Å × 104 Å for SARS-CoV and 94 Å × 144 Å × 100 Å for SAR-CoV-2 systems.^21^

All systems were subjected to equilibration for 50 ps. Following equilibration, 20 ns MD simulations were performed for each system. The velocities and positions of atoms during dynamics were calculated by velocity Verlet integration algorithm^31^ using a time step of 2.0 femtoseconds. All simulations were run with NAMD implementation of Langevin dynamics^32^ at a constant temperature of 298 K using periodic boundary conditions. To model, non-bonding interactions, a switching function was turned on with a ‘switchdist’ of 10 Å, a cutoff of 14 Å, and a ‘pairlistdist’ of 16 Å.

### Binding Free Energy Calculations

Gibbs free energies of binding between the ACE2 and SARS CoV or CoV-2 proteins were calculated using APBS using a standardized method of a treecode-accelerated boundary integral Poisson-Boltzmann equation solver (TABI-PB).^33^ In this method, the protein surface is triangulated, and electrostatic surface potentials are computed. The discretization of surface potentials is utilized to compute the net energy due to solvation as well as electrostatic interactions between the two protein subunits, as outlined in thermodynamic scheme, Scheme 1. Following Scheme 1, the free energy of binding of the two protein fragments in water, can be expressed as a sum of two components (eq. 1)

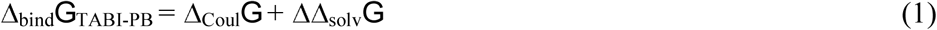

where the Δ_Coul_*G* represents the Coulombic (electrostatic) interactions between the proteins occurring at the protein-protein interface (Scheme 1) and ΔΔ_solv_*G* is the difference of the solvation energies between the complex and the corresponding free proteins:

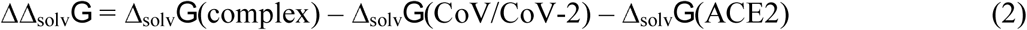

However, the solvation calculation used only part of the entire spike protein as well as the ACE2, therefore Δ_bind_*G*_TABI-PB_ was calibrated by correcting ΔΔ_solv_*G* using experimentally known binding free energy of ACE2…CoV-2:

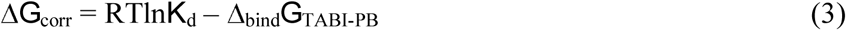

where *K*_d_ is the experimental dissociation constant, which is equal to 37 nM.^16^ The corrected free energy of the solvation

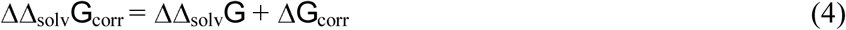

Using the Δ*G*_corr_ and eq. 4, the corrected binding free energy, Δ_bind_*G*^o^ of all protein complexes is expressed by

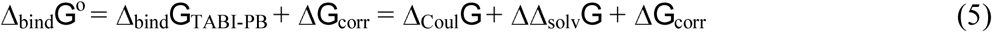

As shown in eq. 4, the combination of last two terms in eq. 5 is equal to ΔΔ_solv_*G*_corr._ Therefore, eq. 5 can be simplified as:

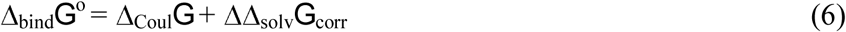

## ACKNOWLEDGEMENTS

We acknowledge computational support from the Blugold Super Computing Cluster (BGSC) of University of Wisconsin-Eau Claire.

## Abbreviations

ACE: Angiotensin Converting Enzyme
CoV: Coronovirus
MD: Molecular Dynamic
RBD: Receptor binding domain
TABI-PB: treecode-accelerated boundary integral Poisson-Boltzmann

---

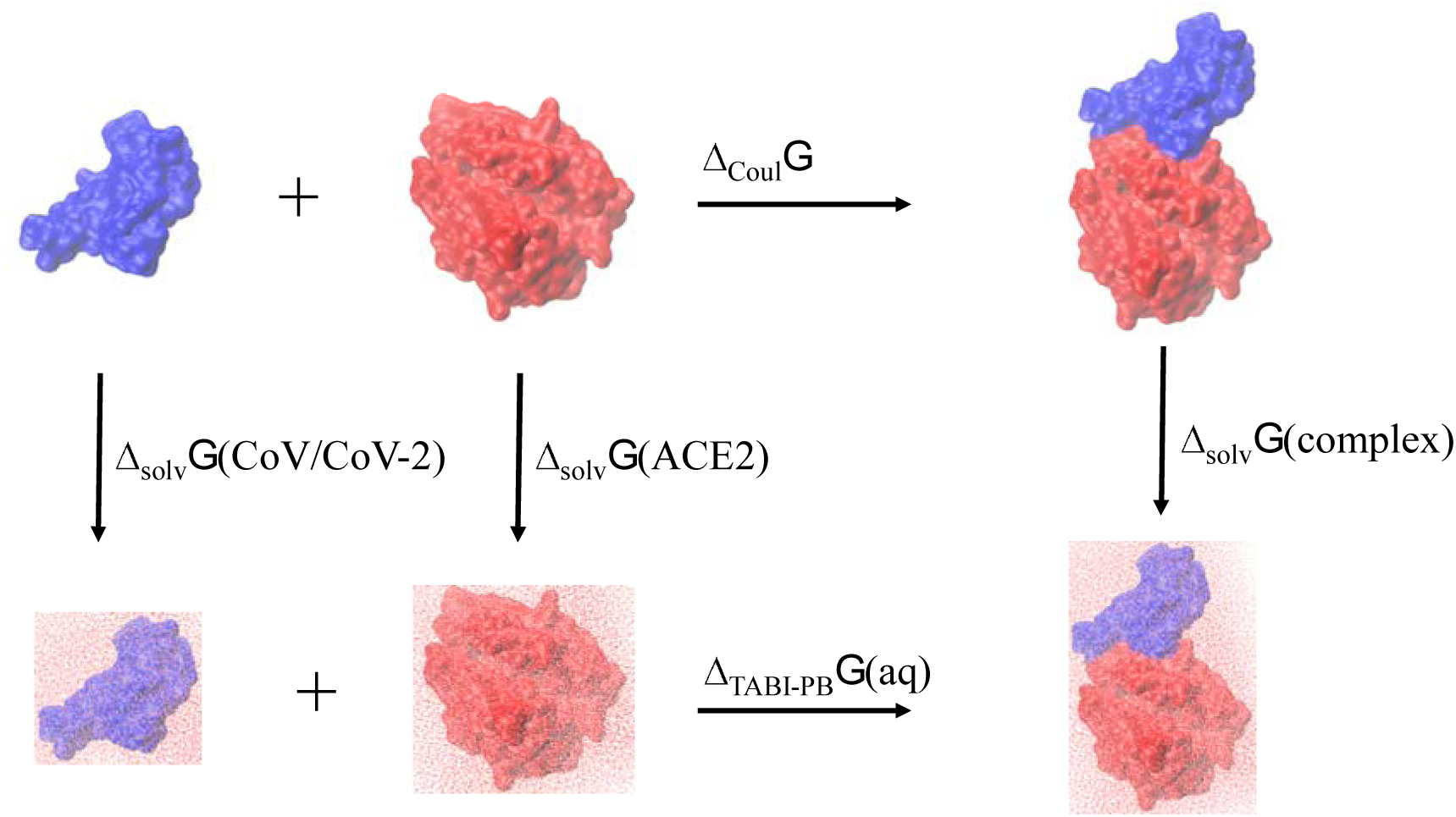
Scheme 1

